# A tradeoff between the ecological and evolutionary stabilities of public goods genes in microbial populations

**DOI:** 10.1101/039784

**Authors:** Joseph Rauch, Jane Kondev, Alvaro Sanchez

**Author notes:** authors for correspondence Joseph Rauch and Alvaro Sanchez.

## Abstract

Microbial populations often rely on the cooperative production of extracellular “public goods” molecules. The cooperative nature of public good production may lead to minimum viable population sizes, below which populations collapse. In addition, “cooperator” public goods producing cells face evolutionary competition from non-producing mutants, or “freeloaders”. Thus, public goods cooperators have to be stable not only to the invasion of freeloaders, but also to ecological perturbations that may push their numbers too small to be sustainable. Through a combination of experiments with microbial populations and mathematical analysis of the Ecological Public Goods Game, we show that game parameters and experimental conditions that improve the evolutionary stability of cooperators also lead to a low ecological stability of the cooperator population. Complex regulatory strategies mimicking those used by microbes in nature may allow cooperators to beat this eco-evolutionary stability tradeoff and become resistant to freeloaders while at the same time maximizing their ecological stability. Our results thus identify the coupled eco-evolutionary stability as being key for the long-term viability of microbial public goods cooperators.

## INTRODUCTION

Populations often require individuals to contribute to their maintenance. Frequently, this involves the production of “public goods”, whose costs are born by the producing individuals but whose benefits are shared by all individuals in the population [1]. Historically, cooperation has been mostly studied as an evolutionary problem. A vast body of work in the field of evolutionary game theory has been devoted to understanding the conditions that lead to the emergence and stability of cooperation, most of which involve some form of self-assortment of cooperators [2–12].

A less appreciated but equally important aspect of social dilemmas is that cooperation may present ecological challenges [13–16]. Populations that require the expression of cooperative traits for their survival, such as the production of a public good, often require large numbers of cooperating individuals in order for the positive effects of their contributions to be significant. When population sizes are too small, the overall production of the public good may not be large enough to sustain the population. This may lead to a minimum viable population size, below which the population collapses [13]. Indeed, the presence of an Allee effect (a positive effect of population size on per capita population growth) in public goods based communities is both predicted by theory [10,11] and has been observed experimentally in microbial populations [13] (Supplementary Fig. 1a).

Ideally thus, a successful public goods cooperator would not only have to be resistant to the evolutionary challenges presented by non-cooperating mutants, but also to ecological perturbations that would push cooperator populations to dangerously low levels (Fig. 1a). Interestingly, both theory [10,11] and experiment indicate that the ecological and evolutionary dynamics of public goods genes may happen at similar timescales, and they are dynamically coupled in a feedback loop [16] (Supplementary Fig. 1b). Given this coupling, it is pertinent to ask whether the stability to evolutionary challenges and the stability to ecological perturbations are also coupled, and whether there exist any constraints that make it easy or difficult to simultaneously maximize the stability to both evolutionary and ecological perturbations.

**Fig. 1:**
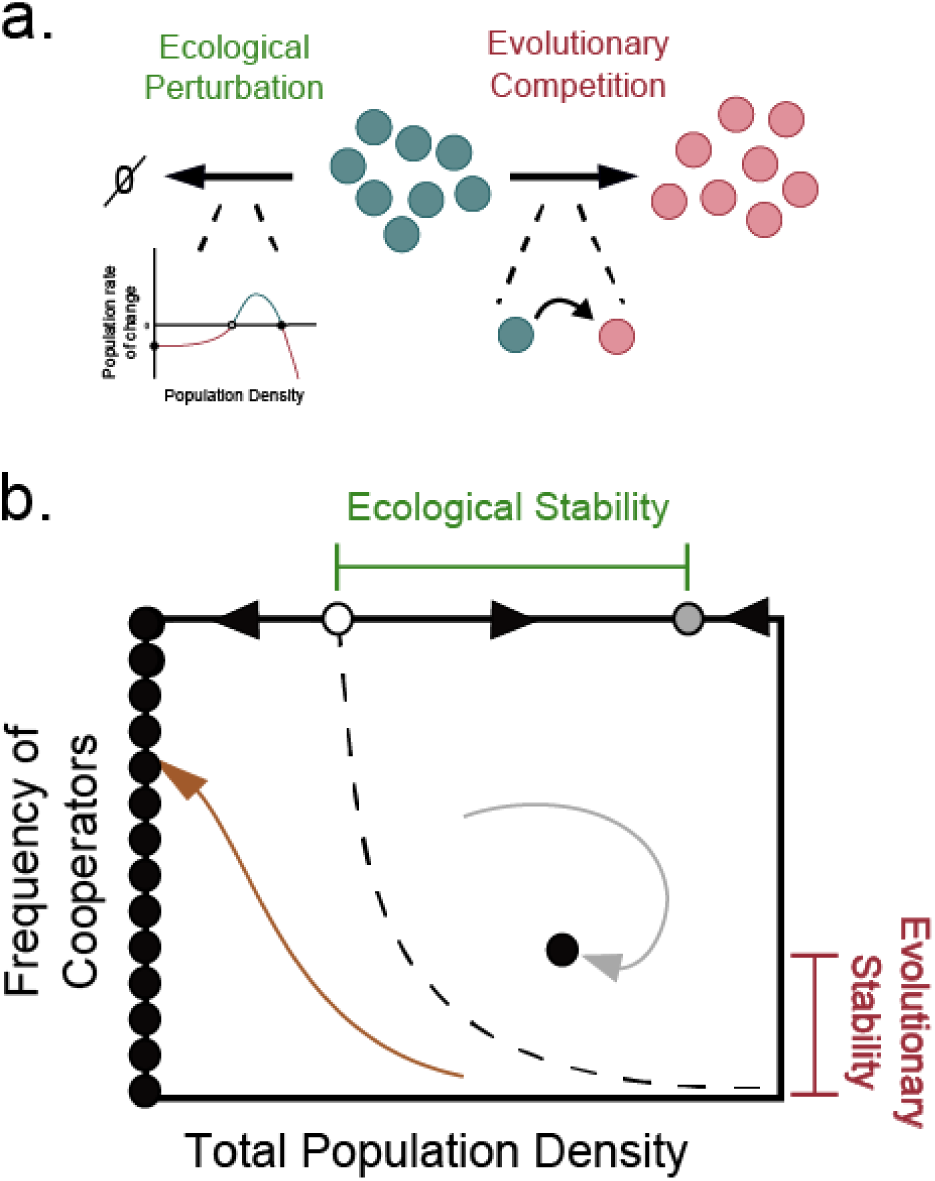
Cooperators face both ecological and evolutionary challenges to their long-term survival. (a) Two different forces can compromise a cooperator allele’s survival. On the one hand freeloaders can emerge by mutation and then take over, driving the cooperator allele to either go extinct or to survive at very low frequencies. Alternatively, the existence of a minimum critical population size may cause external stressors or environmental catastrophes to drive an allele to extinction from purely ecological causes, even in the absence of evolutionary competition with freeloaders. (b) A diagram depicting the measures used for determining the stability of cooperator alleles under both evolutionary and ecological challenges. The evolutionary stability of cooperators to a freeloader invasion is measured as the frequency of cooperators at equilibrium. The ecological stability of a pure population of cooperators (i.e. in the absence of evolutionary competition by freeloaders) is measured as the smallest perturbation to the population size that would cause population collapse. This is the distance between the stable (gray) and unstable (white) fix points in a pure cooperating population.

To investigate this question, we use the cooperative growth of the budding yeast *S. cerevisiae* in sucrose as a model system [17–26]. Budding yeast metabolizes sucrose by secreting an extracellular invertase enzyme that catalyzes sucrose hydrolysis immediately outside of the cell [17,27]. Laboratory yeast strains express this enzyme from a single “public goods” gene, SUC2 [28]. In recent years, sucrose metabolism by yeast cells has emerged as a model system to study public goods cooperation in microbial communities under laboratory conditions [17–26]. It should be noted, however, that it remains unclear whether SUC2 is a true example of public goods cooperation in the wild [29].

## METHODS

### Experimental methods

All experiments were carried out using the same procedures and methods as in reference[16]. Briefly, Strains JG300A and JG210C (*ΔSUC2*) were used as cooperators and freeloaders respectively[15]. JG300A constitutively expresses YFP; JG210C constitutively expresses dTomato. Growth media contained 1X YNB and 1X CSM-his as per manufacturer’s instructions (Sunrise Science), 2% sucrose, 0.001% glucose, and 8 μg/ml histidine. Equal volumes of 200μL of the cell cultures were grown in the 60 internal wells of a flat-bottom 96-well plate (BD Biosciences). To investigate the effects of a chemical stressor on the eco-evolutionary dynamics, 3% ethanol was added to one of the plates. The plates were incubated at either 30°C or 34°C, shaking at 825 rpm. The remaining external wells were filled with 200 μl of growth media and showed no evidence of contamination. The cover of the plate was sealed with parafilm. After a growth period of 23.5hrs, cells were diluted into fresh growth media using different dilution factors (533x, 667x,1333x,1739x,457x,533x, and 800x). The optical density at 620 nm was determined for each well and each plate at the end of every growth cycle, using a Thermo Scientific Multiskan FC microplate spectrophotometer. Frequency of cooperators was determined by flow cytometry using a high throughput Guava cytometer with excitation at 488nm, and detectors at 525nm and 690nm. The frequency of cooperators was established as the number of identified positive hits in the 530nm channel, divided over the sum of hits at both channels.

### Location of the pure cooperator population fixed points

To calculate the stable equilibrium point *(X*_*S*_)we first filtered out the populations that had gone extinct by the last day of the experiment. Then, we averaged the population sizes the surviving populations. To calculate the unstable equilibrium *(X*_*U*_) we found the minimum starting population size that did not go extinct, and the maximum starting population size of those that did go extinct, and calculated the midpoint between the two. The ecological stability of the pure cooperator population was then defined as |log(*X*_*S*_) – log(*X*_*U*_)|.

### Location of the mixed cooperator and freeloader fixed point

To calculate the fixed point for mixed populations of cooperators and freeloaders, we filtered out populations that had gone extinct as described above. We then calculated the relative frequency of cooperators *q(t)* and total population size *u(t)* for each of the surviving populations. We then selected only those populations whose net rate of eco-evolutionary change had slowed down to a slope lower than 0.1, using the criterion: 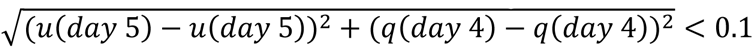 in order to filter out populations which continued to rapidly change and had not yet settled to equilibrium. The population sizes, and relative frequencies of cooperators were then averaged on the last day of the experiment to give the position of the fixed point *(u*_*S*_, *q*_*S*_). The evolutionary stability of the cooperating population is defined as log(*q*_*S*_). Theoretical trajectories for the EPGG, as well as the stable and unstable equilibria were calculated by integrating and analyzing the dynamics equations (see Supplementary Materials) using methods in scipy. Once we knew the positions of all fixed points, we calculated the ecological and evolutionary stability in the same manner as in the laboratory experiment.

## RESULTS

### A tradeoff between ecological and evolutionary stabilities in experimental yeast populations

To probe whether it exists a relationship between ecological stability and the evolutionary stability of SUC2 to freeloader invasion, we grew dozens of yeast cultures in batch mode, with sucrose as the only carbon source and varying environmental parameters such as the dilution factor–akin to a externally imposed death rate-, the growth temperature (30-34C) and the presence and absence of a chemical stressor (3% Ethanol) in the growth media. All of these cultures were started at thirty different initial densities and frequencies of cooperators, and propagated as discussed in the Methods Section (see also Supplementary Fig. 3). After tracking the population size and the frequency of SUC2 for five days, we measured the ecological stability and the evolutionary stability of the SUC2 gene in the population, as described in the Methods section and in Fig. 1b. Then we plotted both stabilities against each other and found that there exists a strong negative correlation between the two: experimental conditions that favor high stability of cooperators to freeloader invasion, also tend to have low ecological stability, and vice versa (Fig. 2a). We also observed a tradeoff between both stabilities when we analyzed two recent independent data sets, where the dilution factor was the only parameter that was altered (Supplementary Fig. 2). Together, these data evidences that a tradeoff between both stabilities is robustly observed in this system.

**Fig. 2:**
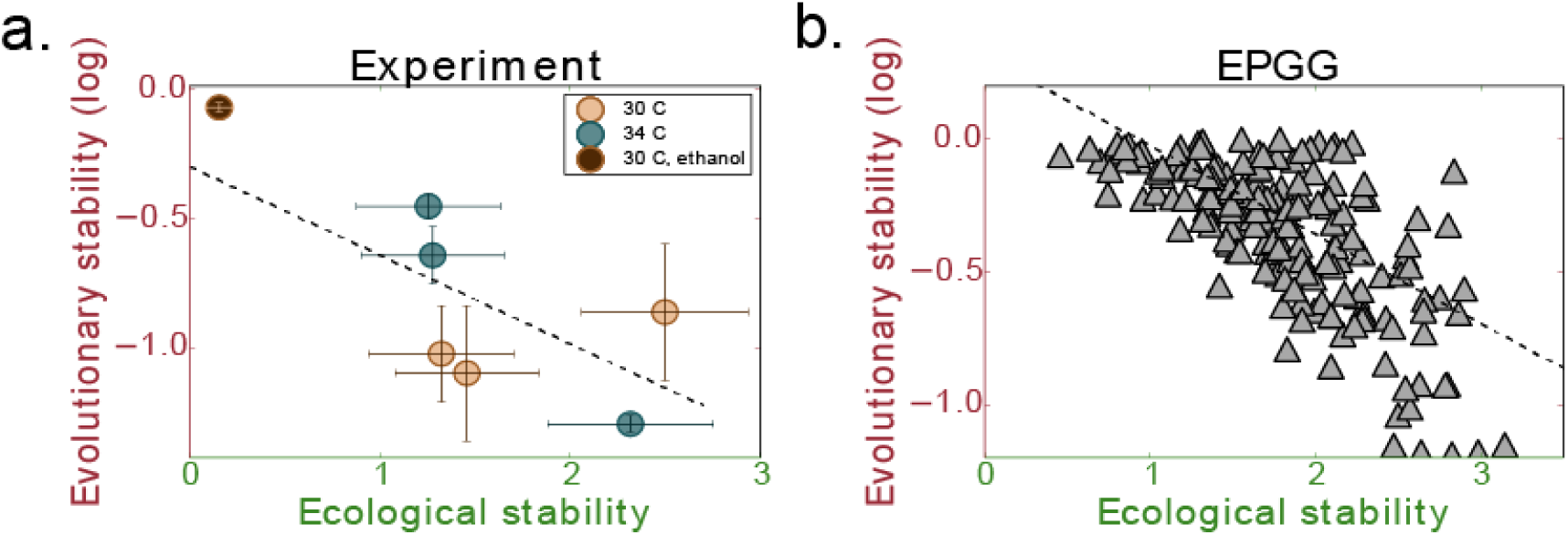
A trade-off between ecological and evolutionary stability is found in the Ecological Public Goods Game and in the yeast-sucrose system. (a) The ecological and evolutionary stability of cooperators measured for yeast growing on sucrose and in several different growth conditions (T=30 °*C* - 34°*C*, [Ethanol]=0%-3%, Dilution Factor=[533x, 667x, 1333x, 1739x, 457x, 533x, and 800x]) for ~30-50 generations. A negative log-log correlation of *ρ* = - 0.61 was found. (b) The ecological and evolutionary stability for cooperators was calculated, as described in Fig. 1b, for 300 different randomly generated parameter sets for the EPGG. The parameters were randomly chosen from the ranges, *r/N* ϵ [0, 1), *N* ϵ [3, 30), *d* ϵ [0.1, 5). All systems had the features in Fig. 1b, with three fix points in the pure cooperator population, and one stable, interior fixed point for mixed populations initialized with a sufficient number of cooperators. The ecological and evolutionary stabilities have a negative log-log correlation *ρ* = - 0.67.

### A tradeoff between ecological and evolutionary stabilities is also observed in theoretical models

To better understand the origin and causes of the observed stability tradeoff, we studied the behavior of the Ecological Public Goods Game (EPGG) [10–12]. Hauert et al [10,11] originally formulated this modeling framework in order to incorporate ecological dynamics to the study of public goods evolution [30–37]. In spite of it being a relatively high level model with no physicochemical parameters, the EPGG model predicted many characteristic features of the eco-evolutionary phase space that were, several years later, observed experimentally in yeast cultures [16] (Supplementary Fig. 1).

In the Public Goods Game, a finite number of *N* players come together to an interaction group. Each player makes an investment of either 1 (if the player is a cooperator), or 0 if it is a freeloader. All of the investments are pooled together, multiplied by a return factor (*r*) and then shared evenly among the *N* participants. The Ecological Public Goods Game is a modification of the Public Goods Game, where the identities of the *N* players are chosen by randomly sampling from a population of cooperators, freeloaders and “vacancies”, whose frequencies are given by *X*, *Y* and *Z* respectively. These “vacancies” are treated as players that neither contribute any investment nor take on any benefit. The payoffs to a focal cooperator or freeloader include the cost of producing the public good and the benefit obtained from their own investment. The payoffs from every possible group composition are averaged and determine the fitness for the cooperators (*ƒ*_*c*_) and freeloaders (*ƒ*_*ƒ*_). A death rate *d* is also assumed for both populations. A more detailed description of the EPGG with its relevant equations is summarized in the Supplementary Text, and described in full in ref. [10,11].

We reasoned that by randomizing the parameters of the game (i.e. the investment return *r*, the size of the interaction group *N*, and the death rate *d*) we could capture the effect of varying different environmental parameters in our yeast cultures, such as the temperature, presence of ethanol and dilution factor. We simulated a total of 300 different eco-evolutionary dynamics using the EPGG model with as many sets of randomized parameters, and for each set the ecological and evolutionary stabilities were determined as discussed in the Methods section and in Fig. 1b. The results are plotted in Fig. 2b, which shows that a tradeoff between both stabilities is also found in the EPGG model.

The model also allows us to understand the existence of this tradeoff. As discussed by Hauert & Doebeli [10–12], small population densities favor cooperators. This is because the benefit that a focal cooperator gets from its own investment gets spread over fewer individuals, and thus it may exceed its cost. In contrast, large population densities lead to fewer vacancies in the interaction groups and thus more other players that will dilute the benefit accrued by a cooperator’s investment; this leads to lower returns from investment and lower cooperator fitness. Hauert and Doebeli also found that the population dynamics of pure cooperator populations are characterized by a strong Allee effect [10,11]: at low population sizes the interaction groups have few cells and low net investments, and therefore small growth rates that may be lower than the death rate (Supplementary Fig. 1). This leads to a minimum “critical” population size below which the net growth rate is negative and populations collapse to extinction. Therefore, game parameters that lead to small population densities at equilibrium will on the one hand benefit cooperators and increase their evolutionary stability against non-cooperator mutant invasion; but on the other hand, these small populations are dangerously close to the critical population size and thus have low ecological stability.

In the theory and experiments discussed above, the strategy of the cooperators (their level of investment in the public good) was kept constant, while the environment changed. It is pertinent to ask then if, for any given environment, an optimal strategy (defined by the level of investment of a player, which may be different from 0 or 1) may exist that simultaneously maximizes the evolutionary and ecological stabilities of cooperators. Given the success of the EPGG model to describe cooperation in microbial populations, we used it to investigate this question.

### Fixed investment strategies may beat the stability tradeoff

We first analyzed the situation when strategies are fixed. In this case, a strategy is defined as the investment level of the cooperators. We start by studying how the equilibrium points and their stability change as we increase or decrease this investment level. We do this by plotting, in Fig. 3a, the bifurcation plot representing the population density of cooperators in equilibrium as a function of the fixed investment level. At low investment levels (red region), the payoffs from the public goods game are too small to support any population. Investments above a critical threshold,

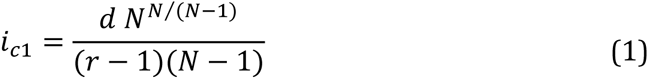

provide enough payoff to sustain cooperator populations, but cannot sustain freeloaders. The stable cooperator population densities are shown by the thick solid line in the blue region of Fig. 3a. For large investments, above the threshold,

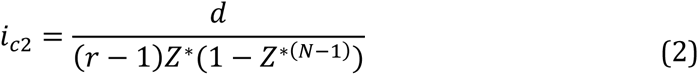

where Z* is the root of the following equation:

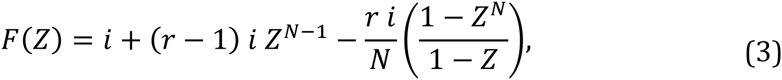

which is the difference between the cooperator and freeloader fitness functions (see Supplementary Text), payoffs from the game are large enough to support a population of both freeloaders and cooperators.

**Fig. 3:**
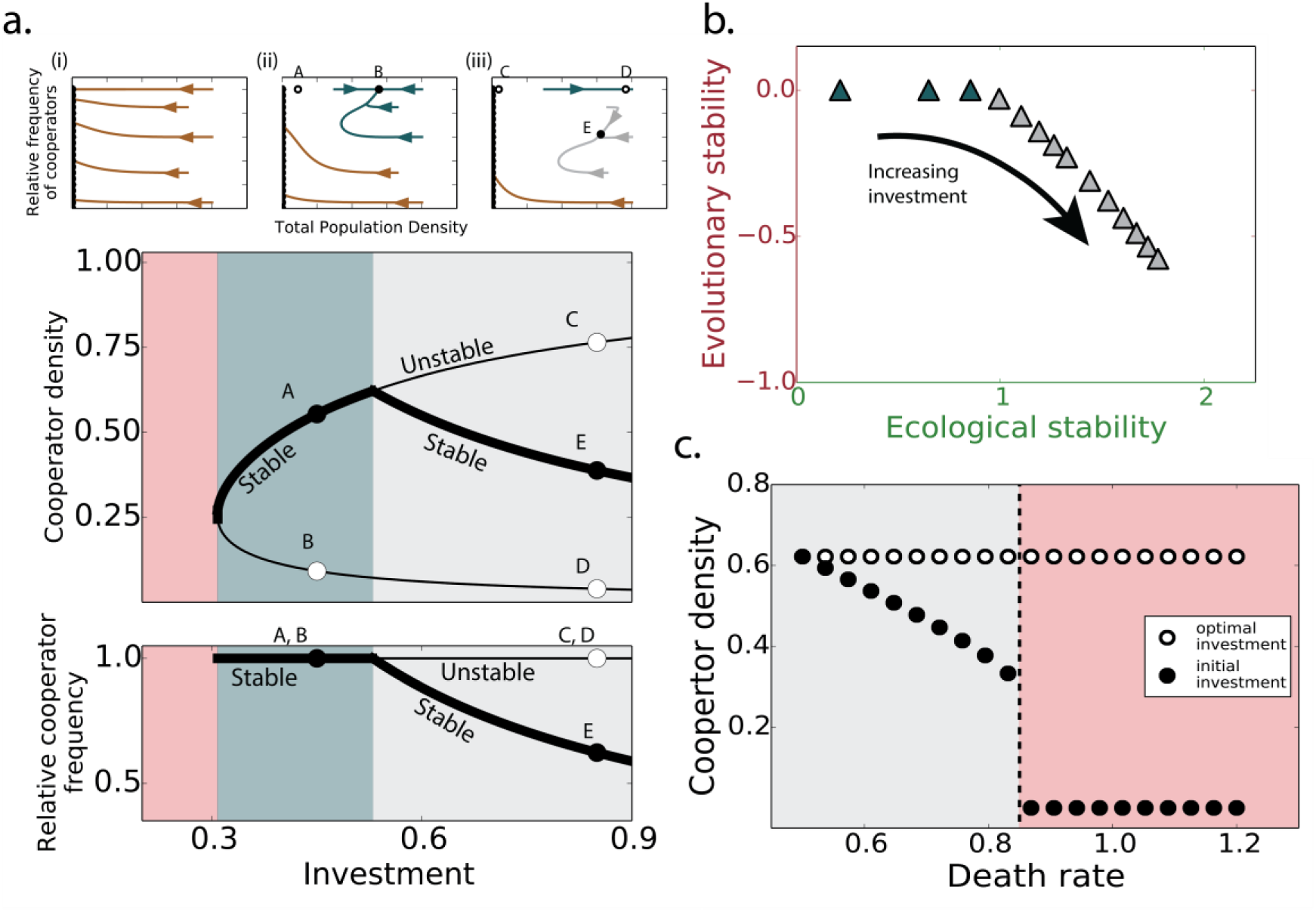
Fixed investment strategies can beat the ecological and evolutionary stability tradeoff within a certain investment range. (a) A bifurcation diagram for the EPGG with unconditional cooperators using the fixed investment as a control parameter. The diagram is split into three regions. Below the critical investment, *i*_*c1*_, cooperators invest too little to support any nonzero population, leading to population collapse (red region, eco-evolutionary phase portrait (i)). Above the critical investment *i*_*c1*_ there is a range of investment levels (blue region, (ii)) for which cooperators are non-invasible by freeloaders. This region ends at the critical investment *i*_*c2*_; above this investment, cooperators can be invaded by freeloaders (gray region, (iii)). The insets (i), (ii), and (iii) are example phase portraits for the red, blue, and gray regions respectively. (b) The evolutionary stability is plotted against the evolutionary stability for increasing investment. (c) An optimal fixed strategy for a particular environment (death rate *d*=0.5) becomes sub-optimal as the environment changes, increasing the death-rate, and it eventually leads to population collapse (experimental observation of this was reported in [13]. For all plots: *r* =5, *N* = 8; for (a, b), *d* = 0.8; and for the insets (i), (ii), and (iii) in (B) respectively; *i* =0.2, *i* = 0.45, *i* = 1.0.

The stable cooperator population in the blue region of Fig. 3a, becomes an unstable saddle point as depicted by its transition to a thin line in the gray region (Fig. 3a), and the emergence of a new stable population of cooperators and freeloaders (Fig. 3a). In this region, as investments increase, the relative frequency of cooperators in the stable population decreases, see Fig. 3a.

To investigate whether by adopting different fixed strategies cooperators may beat the tradeoff, we plot in Fig. 3b the ecological and evolutionary stabilities against each other. Consistent with the findings in Fig 3a, there exist a range of investments for which cooperators are perfectly stable against freeloader invasion, while at the same time their ecological stability may be increased by increasing their investment. At a critical investment (*i*_*c2*_) the tradeoff kicks in and if the investment is increased past that point, the ecological stability continues rising, but the evolutionary stability starts declining. Thus, an optimal strategy may be defined in terms of its maximal evolutionary and ecological stability. Cooperators may beat this tradeoff by adjusting their investment to this optimal level (Fig. 3b).

The value of the optimal investment level depends on the parameters of the game. A cooperator employing a fixed strategy that is optimal in one environment may not even survive at a different environment. For instance, if we modulate the death rate in the model, we find that a strategy that is optimal in one particular set of environmental conditions becomes rapidly suboptimal if the environment changes, and would collapse if the environment deteriorates past a critical point (Fig. 3c).

Although the analysis of fixed strategies is helpful from a conceptual standpoint, the use of purely fixed strategies is not biologically realistic nor it is consistent with the way in which microbes express genes. For instance, the SUC2 gene is conditionally expressed as a function of the accumulation of glucose in the environment. Glucose accumulation is in turn caused by the collective breakdown of sucrose by the cooperator population. Thus, by down-regulating expression of SUC2 yeast cells are able to conditionally adjust their investment in the public good in response to the density of other cooperators around them. We decided to investigate whether conditional (or facultative) cooperation strategies would be advantageous towards beating the stability tradeoff.

### Conditional investment strategies beat the stability tradeoff and are viable over a wider range of environments

Conditional cooperation strategies were modeled using a standard phenomenological model of gene regulatory input-output function known as the Hill Function [38,39]. In particular, we have chosen a Hill function type whose shape is in agreement with the observed input-output function of the SUC2 promoter in *S. cerevisiae* [27], and which takes the form:

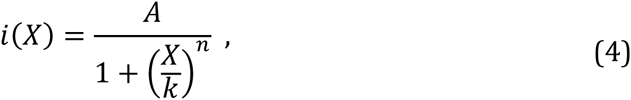

where a cooperator’s investment, *i(X)*, depends on the density of cooperators in the population, *X*. When cooperator frequency *X* is low, *i(X)* is near its maximum (or “amplitude”) *A*; when *X* is equal to the “threshold” *k, i(X)=A/2*; and when cooperator frequency is high, *i(X)* approaches 0. The parameter *n* describes the steepness of the transition from high to low investment. This “Hill function” is a standard model of a gene regulatory input-output function [38] and it matches known patterns of expression of public goods genes by microbes[27,39]. We describe how we modified the EPGG model to account for these conditional strategies in the Supplementary Text.

In order to find how the equilibria and their stabilities change under conditional investment strategies, we need to find the cross-over points between the bifurcation diagram and the investment strategy (Fig. 4a). Those cross-over points mark the investment levels required to keep the population in equilibrium. Different strategies, defined by their threshold (*k*) and amplitude (*A*) values, will cut the bifurcation plot at different places. Therefore, we did not necessarily expect to observe the same dynamic behavior on the phase portrait for different types of strategies. This expectation was confirmed when we analyzed the type of eco-evolutionary phase portrait and the stability of fixed points for different values of *A* and *k* (Fig. 4b). Interestingly this analysis reveals that for large values of the threshold *k*, the type of dynamics on the phase portrait are identical to those previously reported by Hauert and Doebeli for fixed investment levels (gray region). This may explain the success of the fixed investment model to predict the dynamics of yeast populations, in spite of the fact that the expression of the SUC2 gene is not fixed but conditional and follows the function described by equation 4.

**Fig. 4:**
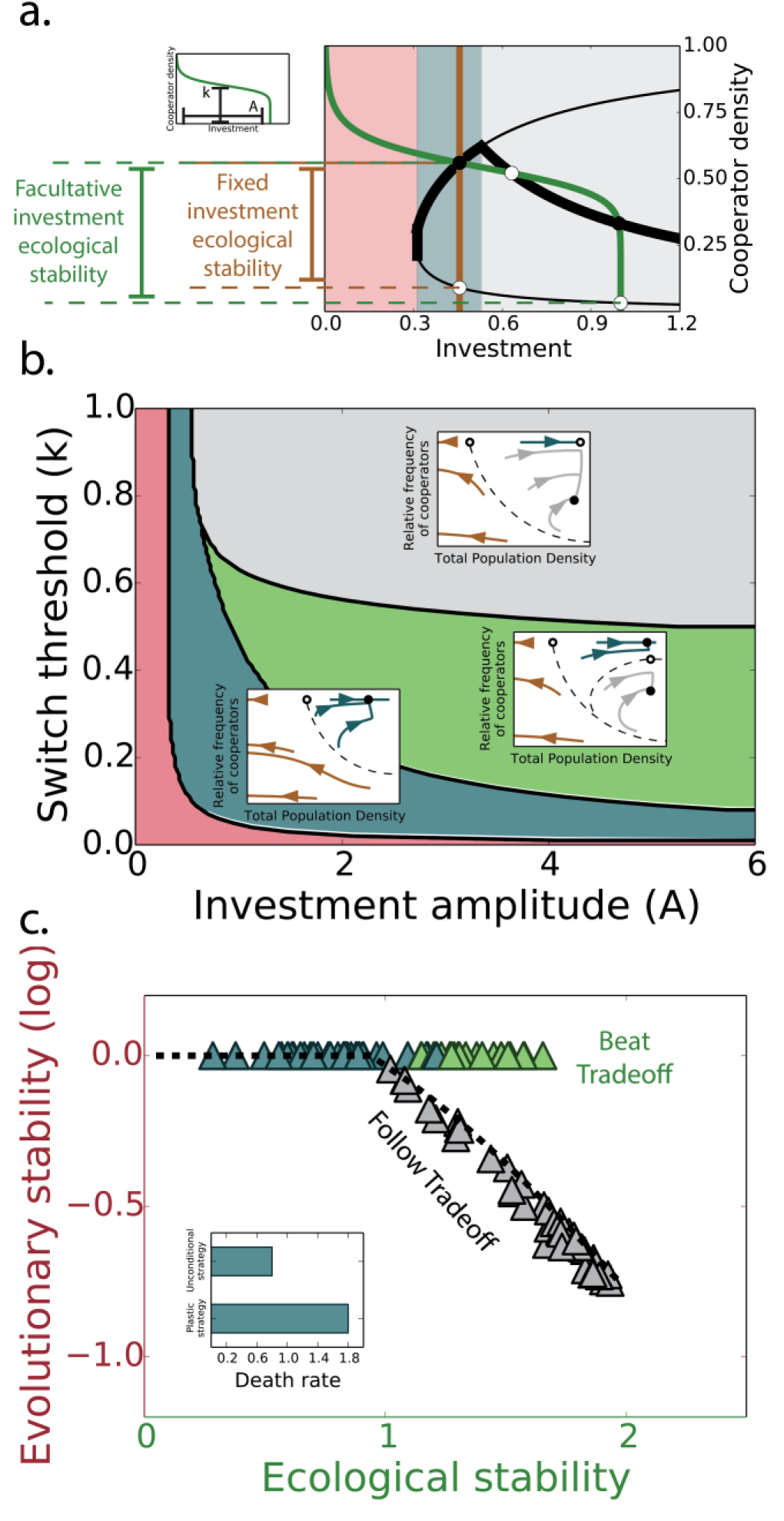
Evolutionary stable strategies (ESS) allow cooperating populations to beat the tradeoff. (a) The same bifurcation diagram depicted in Fig. 3a, with a facultative cooperator’s investment strategy overlaid in green (eq. 4; inset) and a fixed investment strategy overlaid in brown. The intersections between the investment strategies and the bifurcation diagram indicate fixed points. The stability of the fixed points was then analyzed using standard methods and shown by the open (unstable) and closed (stable) circles. The facultative strategy produces two stable and two unstable fixed points aside from a line of stable extinction points and a separatrix which are not shown for clarity. The fixed investment strategy produces two fixed points, one stable and one unstable in addition to the extinction fixed point which is stable. The ecological stability is defined by the distance between the stable and unstable pure cooperator equilibria. This distance is larger for the facultative than for the fixed strategy, as shown. (b) A phase diagram with example phase portraits for different regions of the space of strategies. In the red region, cooperators invest too little and cannot even support themselves. In the blue region, more investment is made and cooperators can support themselves but freeloaders cannot invade. In the green region cooperators are stable but so is the internal, mixed equilibrium point. The gray region leads to phase portraits similar to those observed for unconditional cooperators, with a stable, mixed equilibrium and three pure cooperator equilibria. (c) We plot the evolutionary stability against the ecological stability for 100 different facultative strategies randomly sampled from the strategy space in (b). Points are colored to represent the region of the strategy space to which they belong. Strategies in the gray region follow the same tradeoff reported above for the fixed strategies (dashed line). However, strategies in the blue and green region may beat the tradeoff, and are able to reach very high ecological stabilities without compromising their evolutionary stability, which remains at 1.0. The inset shows how facultative cooperators are capable of surviving harsher conditions. The optimal strategy for both the fixed and facultative cooperators was calculated for one death rate. At other, larger death rates, up to *d* ~ 1.8, facultative cooperators are still capable of surviving while fixed investment cooperators were not. The parameters used in (a) are, *A* = 1.0 and *k* = 0.55 (for the facultative strategy) and *i* = 0.46 (for the fixed investment strategy). The parameters used for the insets in (b) are: *A* = 0.6, *k* = 0.4 for (i), *A* =1.0, *k* = 0.55 for (ii), and *A* = 1.2, *k* = 0.8 for (iii). The parameters used for the inset in (c) are, *A* = 0.7 and *k* = 0.68 for the facultative strategy and *i* = 0.52 for the fixed investment strategy. In all plots: *r* = 5, *N* = 8, and *d* =0.8.

To test whether conditional strategies would be better at beating the stability tradeoff than fixed strategies, we measured the ecological and evolutionary stabilities for 100 random pairs (*A*, *k*) from the space mapped out in Fig. 4b. The results are plotted on Fig. 4c. The solid, brown line on top represents the same results from the fixed strategy. Strikingly, we find that for a range of facultative cooperation strategies (light green region in Fig. 4b), cooperators are able to avoid the tradeoff and achieve high ecological stability while being completely stable against freeloader invasion. This is explained by the location of the intersections between the fixed and facultative strategy lines, and the line of pure cooperator unstable equilibria in the bifurcation diagram (Fig. 4a). As shown in Fig. 4a, the fixed strategy cuts the line of unstable equilibria at a different (and higher) cooperator density than the facultative strategy. This leads to a lower ecological stability for the fixed strategy, even though both strategies have perfect evolutionary stability against mutant freeloaders, and both reach the same population sizes in equilibrium (Fig. 4a).

Furthermore, when we explore the viability of facultative strategies over a range of environmental conditions, we find that they are able to survive (even if they are invasible by freeloaders) at a much wider range of environments, including those for which fixed strategies would go extinct (Fig. 4c-inset). Thus, the overall stability of conditional strategies greatly exceeds that of fixed strategies.

## DISCUSSION

Microbes relying on the production of public goods for their survival face two different types of challenges. On one hand, evolutionary emergence of freeloaders may cause a steep decline in the numbers of cooperative alleles, which code for the expression of the public goods. On the other hand, even when this invasion does not take place, cooperator populations may undergo catastrophic collapses when environmental perturbations push their populations to low levels. Ideally, to ensure their long-term survival, public goods producing microbes should be resistant to both types of challenges.

The work presented in this paper demonstrates that the stabilities to ecological and evolutionary challenges are anti-correlated with each other and exhibit a tradeoff: Environments that favor one diminish the other. This tradeoff is not only observed in laboratory populations of budding yeast growing in sucrose, but more generally they are predicted by the Ecological Public Goods Game theory.

Our analysis suggests that microbes employing facultative cooperation strategies similar to those that have been observed experimentally, may have an enhanced long-term survival as they are able to maximize their evolutionary stability without compromising their ecological stability. This ability to regulate their investment also makes them able to colonize more challenging environments than unconditional cooperators could. Therefore, we establish that in principle there can be a direct relationship between gene regulation and the employment of smart strategies in public goods dilemmas, which may be selected over evolutionary timescales. In future studies, it will be interesting to investigate how other factors, such as multiple cooperating strategies in a community or multiple simultaneous public goods games, alter the eco-evolutionary stability tradeoff.

## Authors’ contributions

JR carried out the modeling work, data analysis, and wrote the paper. JK provided guidance and wrote the paper. AS conceived of the research, designed and performed the experiments, provided guidance, and wrote the paper.

## Acknowledgments

The authors wish to thank Vishal Kottadiel for assisting with one of the daily dilutions as well as members of the Sanchez and Kondev groups for enlightening discussions. The authors also wish to thank Juan Poyatos for his comments on the manuscript and many helpful discussions.

## Supplementary Material

**Supplementary Figures**

**Supplementary Fig. 1:**
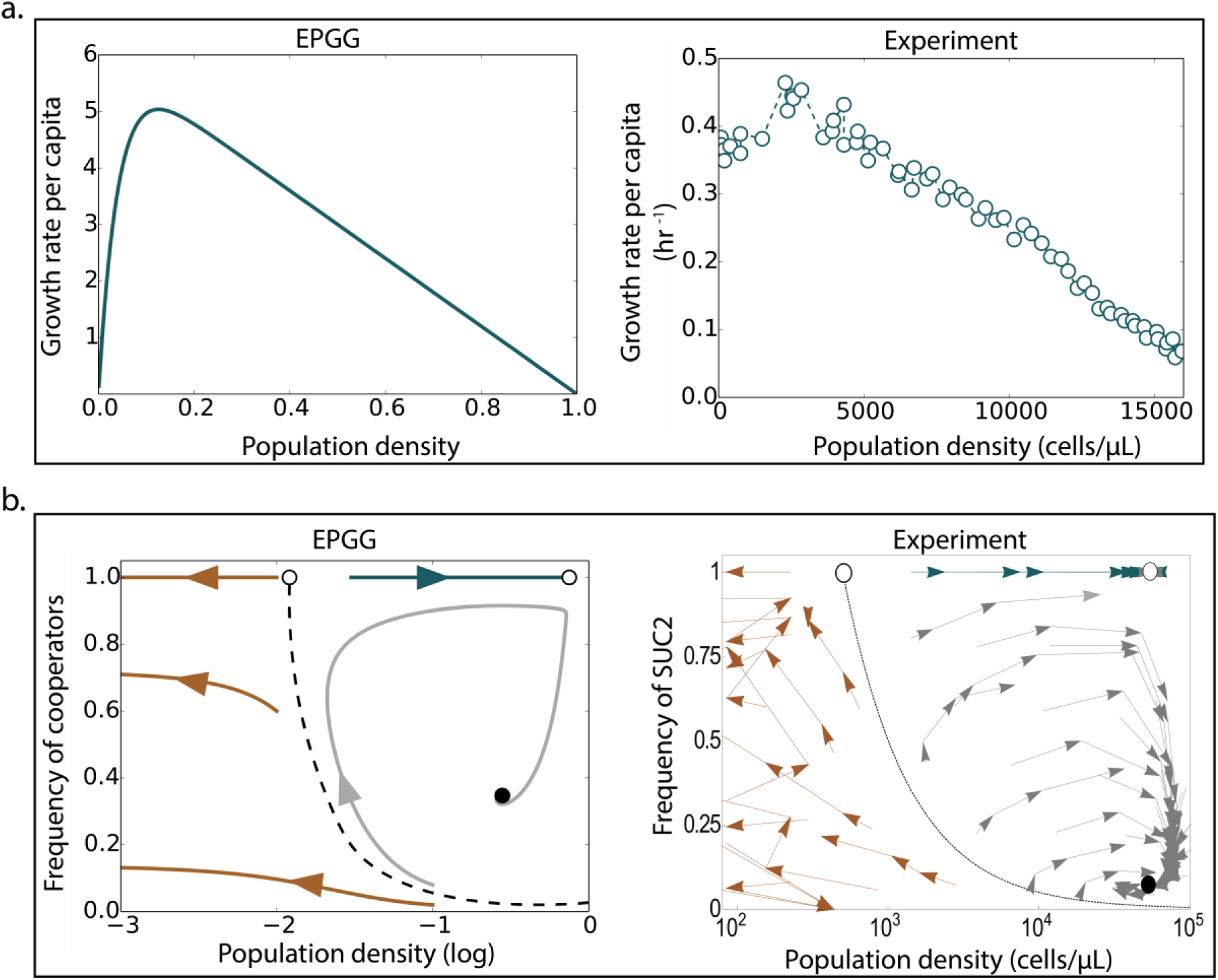
The ecological public goods game (EPGG) predicts many features of the yeast sucrose system. a) A comparison between growth rates predicted by the Ecological Public Goods Game (EPGG) [1,2] on the left, and a reproduced plot from Sanchez and Gore [3] (Copyright 2013 CC-BY) showing experimental growth rates of yeast on the right. In the experiments, cooperators are haploid yeast cells that contain and express the “public good” gene SUC2, which codes for an enzyme that extracellularly breaks down sucrose into glucose and fructose. Freeloaders have a null allele mutant of this gene, and thus enjoy the fructose and glucose produced by the cooperators, without paying the metabolic costs associated to it. Both the prediction by the EPGG and the experimental observations indicate an allee effect, with low growth rates at high and low population densities, and high growth rates at intermediate population densities. b) Theoretical and experimental eco-evolutionary phase portraits for the EPGG [1,2] and yeast experiments [3]. The x-axis, Population Density, reflects population dynamics, whereas the y-axis, frequency of cooperators (SUC2 allele), represents changes in evolutionary dynamics. The yeast experiments observe, and the EPGG model predicts for a wide range of parameters (in this case *r* = 7.0, *N* = 25, *d* = 1.5), the same number and type of fixed points; two unstable saddle points on the pure cooperator line (top boundary), and a stable, mixed population of cooperators and freeloaders. In both phase portraits, a separatrix separates the basins of attraction between the extinct fixed points and the stable, internal fixed point.

**Supplementary Fig. 2:**
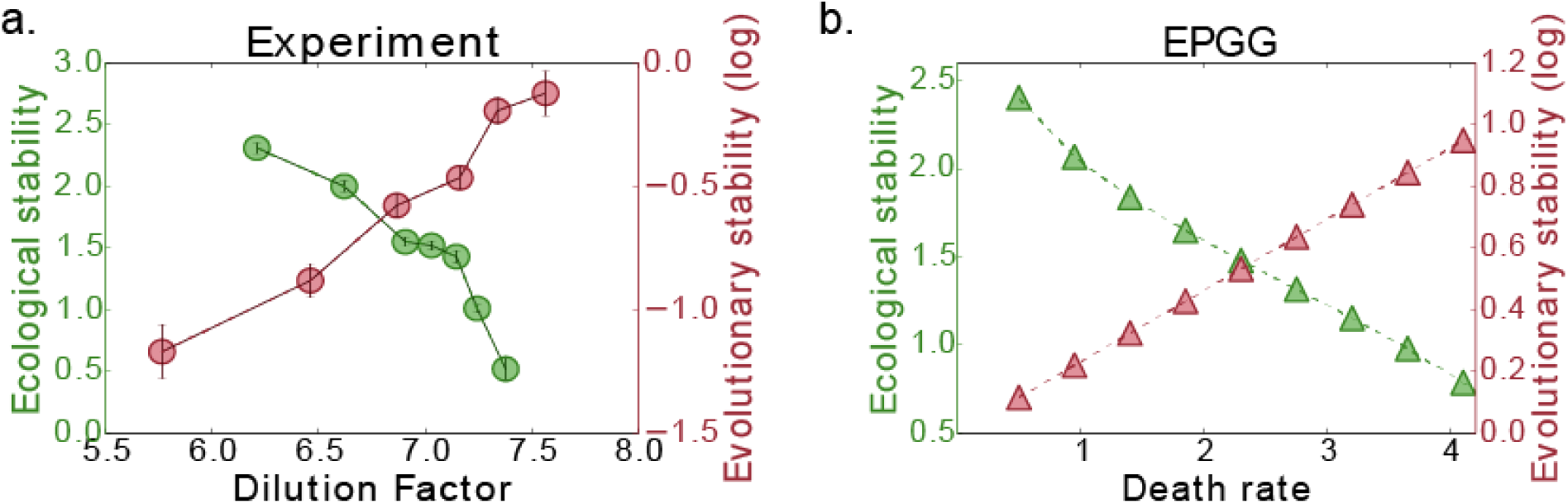
Ecological stability decreases and evolutionary stability increases as the death rate increases in both the yeast-sucrose system and the Ecological Public Goods Game. a) Recently published data of yeast growing in sucrose[4,5] was used to calculate the ecological stability of a pure cooperator population[5] (green)and the evolutionary stability of cooperators to invasion by freeloaders (red) as the dilution factor (an experimentally controllable effective death rate) is varied. As the dilution factor increases the ecological stability diminishes but the evolutionary stability increases. b) Consistent with the yeast experimental data, the EPGG ecological stability decreases as the death rate increase, while the evolutionary stability increases as the death rate increases. Data was simulated by increasing *d* from 0.5 to 5, while fixing *N* = 25, *r* = 7.0.

**Supplementary Fig. 3:**
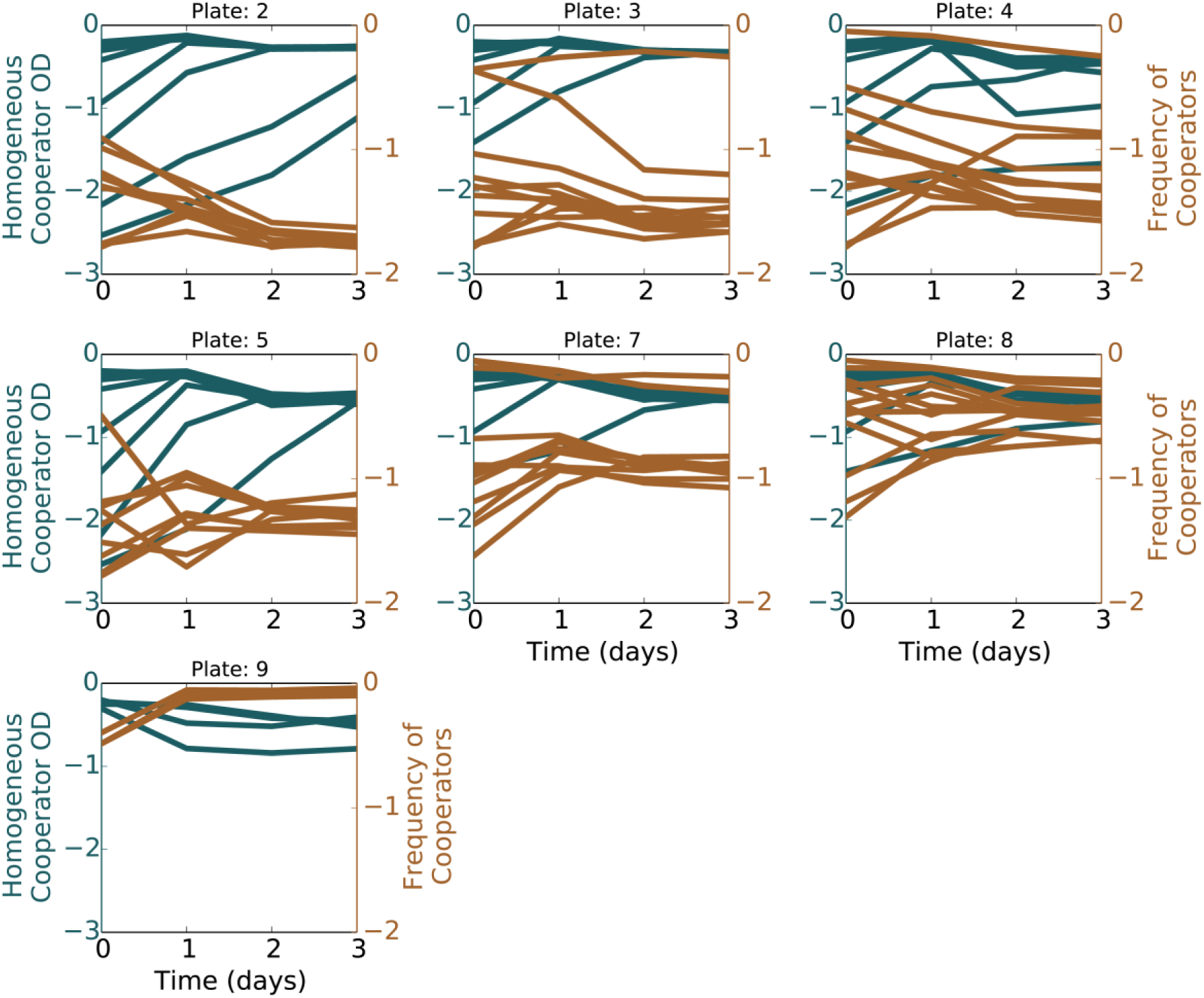
Time traces of yeast populations grown on sucrose under varied conditions. Populations of yeast with and without the SUC gene were competed against each other on sucrose under varied growth conditions for each plate. Blue lines are the population density time traces for pure SUC yeast populations started at different population densities. Most surviving populations reach the same stable equilibrium. Brown lines are SUC frequency time traces for different initial populations of yeast both with and without SUC. Populations were excluded if they had not come to a stable equilibrium, as described in the Methods: Data Analysis section.

## Theory/Mathematical appendix

### Ecological Public Goods Game

Hauert et al [1,2] first developed the Ecological Public Goods Game (EPGG), and it will be summarized here. The EPGG is an iterated public goods game, in which interaction groups for playing the game are constructed by randomly sampling from three populations: cooperators (*X*), freeloaders (*Y*), and ‘vacancies’ (*Z*). Interaction groups have a maximum size, *N*, however, some of those *N* spots can be vacant, neither contributing nor benefiting from the game. This couples the payoffs from the public goods game to the population dynamics, as the total number of players splitting the public good will vary as the as the total population density, *u* = *X* + *Y*, changes. The populations change according to:

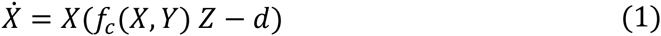

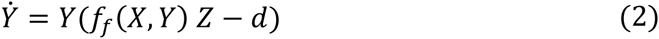

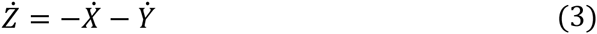

The las equation comes from the constraint on the populations, 1 = *X* + *Y* + *Z*, which limits the size of the total population. The fitness functions, *f*_*c*_ and *f*_*f*_, are determined by the payoffs from the public goods game, and determine the growth of each population. They are scaled by *Z*, the amount of vacant space in the system, to determine the per capita growth rate for each population. This requires that populations only grow into available, vacant space. A per capita death rate, *d*, is also imposed on the populations.

### Derivation of fitness functions

The traditional Public Goods Game is played as follows. Each cooperator in the interaction group makes an investment, *i*, into the production of the public good, while freeloaders invest nothing. The total investment is multiplied by a return factor, *r*, and then distributed amongst all participants, cooperator and freeloader alike. The payoffs for an individual freeloader, *P*_*f*_, or cooperator, *P*_*c*_, in a group of *m* other cooperators and *G* total players look like:

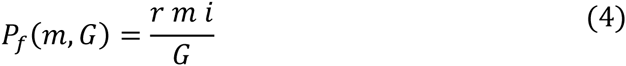

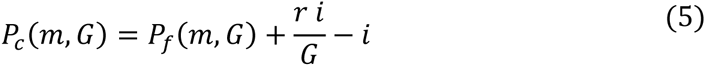

The cooperator receives an extra payout, *r·i/G*, from its own investment, for which it pays *i*. In the EPGG, the interaction group can also be composed of vacancies, not just cooperators and freeloaders. This implied that when forming interaction groups via random sampling, not only will the number of cooperators vary group to group, but also the total number of players, *G*. The fitness functions are determined by averaging the payoffs from every possible interaction group capable of being formed by randomly sampling from the population of cooperators, freeloaders, and ‘vacancies’. The average is weighted by the probability of each interaction group forming, using the normalized population densities, *X*, *Y*, and *Z*, and are formed with replacement, implying infinite population (where 1=*X*+ *Y*+*Z* constrains the density of the populations).

To average payoffs, we first average over the possible number of cooperators within a group holding the group size (*G* the number of cooperators and freeloaders only) constant. The probability of finding *m* cooperators in a group of size *G* is binomially distributed, as *G* only counts cooperators and freeloaders. Thus, the average payoff for a freeloading or cooperating individual in a group of size *G* is:

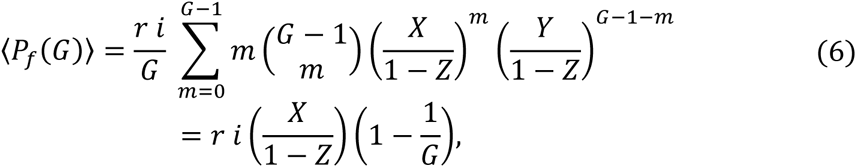

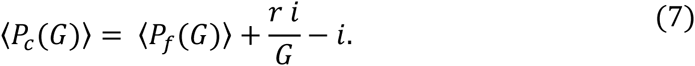

The probabilities for interacting with a cooperator or freeloader are renormalized by dividing by (1-*Z*), so as to determine the relative frequency of each population. Randomly formed groups may not only vary in cooperator count, *m*, but also group size, *G*. Thus, after averaging over the variation in cooperator count between groups of the same size, the payoffs are averaged again, this time over groups of different sizes. The probability of being in a group of size *G* is again binomially distributed, with the probability of interacting with another player (cooperator or freeloader) being *1-Z*, and the probability of not meeting a player being *Z*. The result is the average per capita payoff within each population, the per capita fitness:

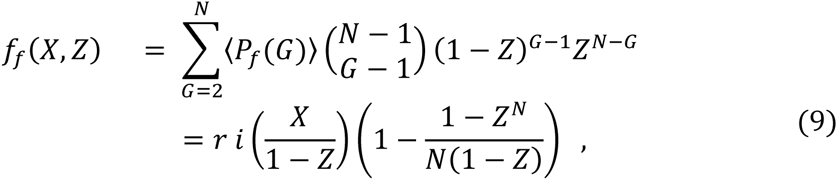

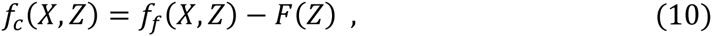

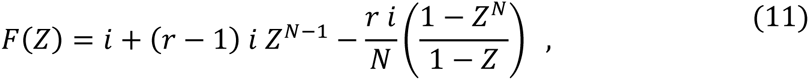

where *f*_*f*_ is the fitness function for the freeloading population, *f*_*c*_ is the fitness function for the cooperating population, and the function *F(Z)* is the difference between the freeloading and cooperating fitness. With these fitness equations, the system defined by equations 1, 2, and 3 is complete. However, the eco-evolutionary dynamics can be made more apparent with a change of variables. Using the relative fraction of cooperators, *q = X/(1-Z)*, to describe evolutionary dynamics between the cooperators and freeloaders, and the total population density, *u = 1-Z*, to describe the ecological dynamics affecting the total population, the new dynamical equations are:

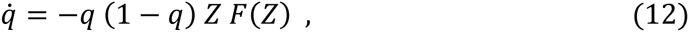

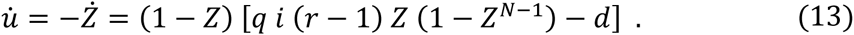

The dynamical system in eqs. 12 and 13 can be more useful for analyzing the eco-evolutionary dynamics of the EPGG.

### Including plastic investment strategies

In the model above, cooperators make an unconditional investment, *i*, into the production of a public good. However, one can imagine another strategy, where a cooperators investment changes depending on the environment the cooperator is experiencing. Inspired by the expression profile of many different genes, particularly the SUC2 gene in yeast, we investigated investment strategies of the form:

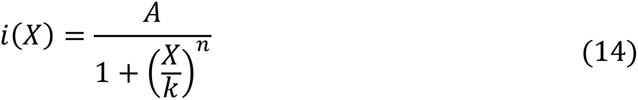

Where *X* is the cooperator density, with *A* being the maximum investment at *X*=0, and *k* determines the density at which cooperators invest half the maximum. The rapidity cooperators switch between high and low levels of investment is determined by *n*. We understand this strategy to mean a cooperator in a cooperator scarce environment, will invest as much as it can to the public good in an effort to survive. However, when cooperators are dense, the environment is already rich in the public good and only a small investment is needed to sustain the population. With an investment based only on cooperator density, it is also easy to incorporate into the model defined by equations 12 and 13 by substituting *i* for *i(X)*, where *X*= *u·q*.

### Analyzing the EPGG

As Hauert et al point out in [1,2], a homogeneous population of unconditional cooperators is capable of surviving at a stable nonzero population density, as long as 1<[*i*(*r*-1)(*N*-1)]/[*d N*^-*N/*(*N*-1)^]. By rearranging this inequality, we can find a minimum investment required for populations to survive,

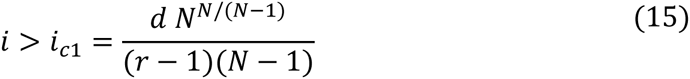

Investments below *i*_*c1*_ are too low to sustain even a homogeneous cooperating population. We can also calculate the investment level at which a freeloader can favorably mutate into a homogeneous cooperating population. We use equation 13 to find the investment, *i*_*c2*_, at which *q\**=1 and *u\** is the root of the difference function, found by setting equation 12 equal to zero,

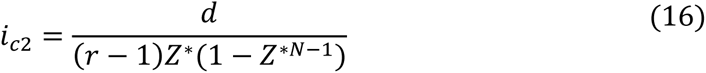

Investments below *i*_*c1*_ are too low to support any population. Cooperators making investment above *i*_*c1*_ but below *i*_*c2*_ can sustain a stable population which is unfavorable to freeloader mutants. Finally, investments above *i*_*c2*_ are large enough to support stable populations of cooperators and freeloader, giving more and more an evolutionary advantage to freeloaders as the investment in increased further.

## References

1. Levin, S. A. 2014 Public goods in relation to competition, cooperation, and spite. Proc. Natl. Acad. Sci. U. S. A. 111, 10838–10845. (doi:10.1073/pnas.1400830111)

2. Nowak, M. A. 2006 Five rules for the evolution of cooperation. Science 314, 1560–1563. (doi:10.1126/science.1133755)

3. Hamilton, W. D. 1964 The genetical evolution of social behaviour. I. J. Theor. Biol. 7, 1–16. (doi:10.1016/0022-5193(64)90038-4)

4. Axelrod, R. & Hamilton, W. D. 1981 The evolution of cooperation. Science 211, 1390–1396.

5. West, S. A., Griffin, A. S. & Gardner, A. 2007 Social semantics: altruism, cooperation, mutualism, strong reciprocity and group selection. J. Evol. Biol. 20, 415–432. (doi:10.1111/j.1420-9101.2006.01258.x)

6. Ohtsuki, H., Hauert, C., Lieberman, E. & Nowak, M. A. 2006 A simple rule for the evolution of cooperation on graphs and social networks. Nature 441, 502–505. (doi:10.1038/nature04605)

7. Traulsen, A. & Nowak, M. A. 2006 Evolution of cooperation by multilevel selection. Proc. Natl. Acad. Sci. 103, 10952–10955. (doi:10.1073/pnas.0602530103)

8. Sachs, J. L., Mueller, U. G., Wilcox, T. P. & Bull, J. J. 2004 The Evolution of Cooperation. Q. Rev. Biol. 79, 135–160. (doi:10.1086/383541)

9. Axelrod, R. M. 2006 The evolution of cooperation. New York: Basic Books.

10. Hauert, C., Wakano, J. Y. & Doebeli, M. 2008 Ecological public goods games: Cooperation and bifurcation. Theor. Popul. Biol. 73, 257–263. (doi:10.1016/j.tpb.2007.11.007)

11. Hauert, C., Holmes, M. & Doebeli, M. 2006 Evolutionary games and population dynamics: maintenance of cooperation in public goods games. Proc. R. Soc. B Biol. Sci. 273, 2565–2571.

12. Wakano, J. Y., Nowak, M. A. & Hauert, C. 2009 Spatial dynamics of ecological public goods. Proc. Natl. Acad. Sci. 106, 7910–7914. (doi:10.1073/pnas.0812644106)

13. Dai, L., Vorselen, D., Korolev, K. S. & Gore, J. 2012 Generic Indicators for Loss of Resilience Before a Tipping Point Leading to Population Collapse. Science 336, 1175–1177. (doi:10.1126/science.1219805)

14. Chen, A., Sanchez, A., Dai, L. & Gore, J. 2014 Dynamics of a producer-freeloader ecosystem on the brink of collapse. Nat. Commun. 5. (doi:10.1038/ncomms4713)

15. Harrington, K. I. & Sanchez, A. 2014 Eco-evolutionary dynamics of complex social strategies in microbial communities. Commun. Integr. Biol. 7. (doi:10.4161/cib.28230)

16. Sanchez, A. & Gore, J. 2013 Feedback between population and evolutionary dynamics determines the fate of social microbial populations. PLoS Biol. 11, e1001547.

17. Greig, D. & Travisano, M. 2004 The Prisoner’s Dilemma and polymorphism in yeast SUC genes. Proc. R. Soc. Lond. B Biol. Sci. 271, S25–S26. (doi:10.1098/rsbl.2003.0083)

18. Dai, L., Korolev, K. S. & Gore, J. 2015 Relation between stability and resilience determines the performance of early warning signals under different environmental drivers. Proc. Natl. Acad. Sci. U. S. A. 112, 10056–10061. (doi:10.1073/pnas.1418415112)

19. Datta, M. S., Korolev, K. S., Cvijovic, I., Dudley, C. & Gore, J. 2013 Range expansion promotes cooperation in an experimental microbial metapopulation. Proc. Natl. Acad. Sci. U. S. A. 110, 7354–7359. (doi:10.1073/pnas.1217517110)

20. Dai, L., Korolev, K. S. & Gore, J. 2013 Slower recovery in space before collapse of connected populations. Nature 496, 355–358. (doi:10.1038/nature12071)

21. Craig Maclean, R. & Brandon, C. 2008 Stable public goods cooperation and dynamic social interactions in yeast. J. Evol. Biol. 21, 1836–1843. (doi:10.1111/j.1420-9101.2008.01579.x)

22. Celiker, H. & Gore, J. 2012 Competition between species can stabilize public-goods cooperation within a species. Mol. Syst. Biol. 8, 621. (doi:10.1038/msb.2012.54)

23. MaClean, R. C., Fuentes-Hernandez, A., Greig, D., Hurst, L. D. & Gudelj, I. 2010 A mixture of ‘cheats’ and ‘co-operators’ can enable maximal group benefit. PLoS Biol. 8. (doi:10.1371/journal.pbio.1000486)

24. Van Dyken, J. D., Müller, M. J. I., Mack, K. M. L. & Desai, M. M. 2013 Spatial population expansion promotes the evolution of cooperation in an experimental Prisoner’s Dilemma. Curr. Biol. CB 23, 919–923. (doi:10.1016/j.cub.2013.04.026)

25. Koschwanez, J. H., Foster, K. R. & Murray, A. W. 2011 Sucrose utilization in budding yeast as a model for the origin of undifferentiated multicellularity. PLoS Biol. 9, e1001122. (doi:10.1371/journal.pbio.1001122)

26. Koschwanez, J. H., Foster, K. R. & Murray, A. W. 2013 Improved use of a public good selects for the evolution of undifferentiated multicellularity. eLife 2, e00367. (doi:10.7554/eLife.00367)

27. Gore, J., Youk, H. & van Oudenaarden, A. 2009 Snowdrift game dynamics and facultative cheating in yeast. Nature 459, 253–256. (doi:10.1038/nature07921)

28. Carlson, M., Osmond, B. C. & Botstein, D. 1981 Mutants of Yeast Defective in Sucrose Utilization. Genetics 98, 25–40.

29. Bozdag, G. O. & Greig, D. 2014 The genetics of a putative social trait in natural populations of yeast. Mol. Ecol. 23, 5061–5071. (doi:10.1111/mec.12904)

30. Brown, S. P. & Taddei, F. 2007 The durability of public goods changes the dynamics and nature of social dilemmas. PloS One 2, e593. (doi:10.1371/journal.pone.0000593)

31. Sasaki, T., Okada, I. & Unemi, T. 2007 Probabilistic participation in public goods games. Proc. Biol. Sci. 274, 2639–2642. (doi:10.1098/rspb.2007.0673)

32. Parvinen, K. 2010 Adaptive dynamics of cooperation may prevent the coexistence of defectors and cooperators and even cause extinction. Proc. R. Soc. B Biol. Sci., rspb20100191. (doi:10.1098/rspb.2010.0191)

33. Korolev, K. S. 2013 The fate of cooperation during range expansions. PLoS Comput. Biol. 9, e1002994. (doi:10.1371/journal.pcbi.1002994)

34. Zhang, F. & Hui, C. 2011 Eco-evolutionary feedback and the invasion of cooperation in prisoner’s dilemma games. PloS One 6, e27523. (doi:10.1371/journal.pone.0027523)

35. Allen, B. & Nowak, M. A. 2013 Cooperation and the Fate of Microbial Societies. PLoS Biol. 11. (doi:10.1371/journal.pbio.1001549)

36. Huang, W., Hauert, C. & Traulsen, A. 2015 Stochastic game dynamics under demographic fluctuations. Proc. Natl. Acad. Sci. U. S. A. 112, 9064–9069. (doi:10.1073/pnas.1418745112)

37. Waite, A. J., Cannistra, C. & Shou, W. 2015 Defectors Can Create Conditions That Rescue Cooperation. PLoS Comput. Biol. 11, e1004645. (doi:10.1371/journal.pcbi.1004645)

38. Frank, S. A. 2013 Input-output relations in biological systems: measurement, information and the Hill equation. Biol. Direct 8, 31. (doi:10.1186/1745-6150-8-31)

39. Kuhlman, T., Zhang, Z., Saier, M. H. & Hwa, T. 2007 Combinatorial transcriptional control of the lactose operon of Escherichia coli. Proc. Natl. Acad. Sci. 104, 6043–6048. (doi:10.1073/pnas.0606717104)

